# Genomic and transcriptomic signals of thermal tolerance in heat-tolerant corals (*Platygyra daedalea*) of the Arabian/Persian Gulf

**DOI:** 10.1101/185579

**Authors:** Nathan L. Kirk, Emily J. Howells, David Abrego, John A. Burt, Eli Meyer

**Affiliations:** Department of Integrative Biology, Oregon State University. Corvallis, OR 97331; Marine Biology Laboratory, CGSB, New York University Abu Dhabi, Abu Dhabi, United Arab Emirates.; Department of Natural Science and Public Health, Zayed University, Abu Dhabi, United Arab Emirates.

**Keywords:** Climate adaptation, heritability, transcriptional stress response

## Abstract

Scleractinian corals occur in tropical regions near their upper thermal limits, and are severely threatened by rising ocean temperatures. Ocean warming leads to loss of symbiotic algae (*Symbiodinium*), reduced fitness for the coral host, and degradation of the reef. However, several recent studies have shown that natural populations of corals harbor genetic variation in thermal tolerance that may support adaptive responses to warming. Here we’ve extended these approaches to study heat tolerance of corals in the Persian/Arabian Gulf, where heat-tolerant local populations have adapted to warm summer temperatures (>36°C). To evaluate whether selection has depleted genetic variation in thermal tolerance, estimate the potential for future adaptive responses, and understand the functional basis for these corals’ unusual heat tolerance, we measured thermal tolerance using controlled crosses in the Gulf coral *Platygyra daedalea*. We found that heat tolerance is highly heritable in this population (0.487-0.748), suggesting substantial potential for adaptive responses to selection for thermal tolerance. To identify genetic markers associated with this variation, we conducted genomewide SNP genotyping in parental corals and tested for relationships between paternal genotype and thermal tolerance of the offspring. We found that multilocus SNP genotypes explained a large fraction of variation in thermal tolerance in these crosses (69%). To investigate the functional basis of these differences in thermal tolerance, we profiled transcriptional responses in tolerant and susceptible families, revealing substantial sire effects on transcriptional responses to thermal stress. We also studied sequence variation in these expressed sequences, identifying alleles and functional groups associated with thermal tolerance. Our findings demonstrate that corals in these populations harbor extensive genetic variation in thermal tolerance, and these heat-tolerant phenotypes differ in both gene sequences and transcriptional stress responses from their susceptible counterparts.

## Introduction

Global climate change is rapidly changing conditions and warming the globe as the atmospheric concentrations of CO_2_ increase. This change affects many organisms that now must cope with altered environments. In response to warming, organisms can migrate either as adults or larvae, adapt or acclimatize to novel conditions, or simply perish via extirpation or even extinction (O’Connor *et al.*, 2012). Despite only ~1° C increase in global temperature since preindustrial times, there have been numerous changes observed in populations and communities including modified behaviors, morphologies, physiologies, distributions, phenologies, interspecific relations and even changes in community productivities (reviewed in Scheffers *et al.*, 2016). Not all organisms will be equally affected with some increasing their distribution and/or abundances across taxonomic groups (Harvey *et al.*, 2013, Scheffers *et al.*, 2016).

Atmospheric CO_2_ levels and concomitant warming also affects the oceans, which act as a large sink for anthropogenic emissions reshaping biogeochemical cycles and consequently marine life. How global climate change will affect marine organisms remains relatively understudied compared to terrestrial systems despite the ecosystem services provided by oceans and increasing evidence that marine species are impacted at faster temporal scales (Gattuso *et al.*, 2015, Hoegh-Guldberg & Bruno, 2010, Richardson *et al.*, 2012). Global oceanic temperatures have been rising and are predicted to rise further over the next century with or without mitigation steps highlighted in the recent Copenhagen accord (Gattuso *et al.*, 2015, Harvey *et al.*, 2013). Warming is accompanied by increased acidification, which can act synergistically to negatively impact calcifying species (Harvey *et al.*, 2013, Hoegh-Guldberg *et al.*, 2007). Although many marine taxa will be impacted by climate change, scleractinian corals are often hypothesized to be one of the most imperiled clades (Doney *et al.*, 2012, Gattuso *et al.*, 2015, Hoegh-Guldberg *et al.*, 2007, van Oppen *et al.*, 2017, Veron *et al.*, 2009).

Coral survival will depend on the ability to acclimatize and adapt to increasing water temperatures. Some adult colonies can gain some degree of thermal protection from their algal symbionts and a growing literature demonstrates that harboring different species and strains of symbiotic dinoflagellates correlates with increased host tolerance (Berkelmans & van Oppen, 2006, LaJeunesse *et al.*, 2014, Silverstein *et al.*, 2015). The symbiotic algal complement has been hypothesized to be the driving force of coral acclimatization and quick adaptation to warming waters (Baker, 2004, Buddemeier *et al.*, 2004). More recently, the role of the coral itself in acclimatization and adaptation is becoming more appreciated (Baird *et al.*, 2009, Barshis *et al.*, 2013, Bay & Palumbi, 2014, Bellantuono *et al.*, 2012b, Kenkel & Matz, 2016, Logan *et al.*, 2014, Palumbi *et al.*, 2014, Putnam & Gates, 2015). Corals with different thermal histories have differential mortality and bleaching susceptibilities (Dixon *et al.*, 2015, Maynard *et al.*, 2008, Middlebrook *et al.*, 2008). Corals from populations exposed to locally warmer and also more variable temperatures generally have higher bleaching thresholds, lower mortality, and greater gene expression plasticity compared to corals from nearby cooler and stable areas (Oliver & Palumbi, 2011, Schoepf *et al.*, 2015, Thompson & van Woesik, 2009, van Woesik *et al.*, 2012). A growing number of studies have demonstrated transcriptional changes associated with heat stress (Barshis *et al.*, 2013, Bay & Palumbi, 2014, Bellantuono *et al.*, 2012a, DeSalvo *et al.*, 2008, Kaniewska *et al.*, 2015, Kenkel & Matz, 2016, Meyer *et al.*, 2011, Palumbi *et al.*, 2014, Rosic *et al.*, 2014, Voolstra *et al.*, 2009) and tolerant corals are able to respond in part by having altered transcriptional profile relative to susceptible colonies (Barshis *et al.*, 2013, Bay & Palumbi, 2014, Kenkel & Matz, 2016, Palumbi *et al.*, 2014). Likewise, thermal protection can be passed to future generations by acclimation of parental colonies (Putnam & Gates, 2015). There are corals that naturally exist in warm waters that consistently reach temperatures above the thermal maximum on other reefs (e.g. Krueger *et al.*, 2017). Corals living in these warm environments can be used as a proxy for future climate change scenarios and to help further understand the role of acclimatization and adaptation in coral survival.

One of the warmer regions with coral reefs is the Arabian/Persian Gulf (hereafter referred to as the Gulf) where wet-bulb air temperatures, which are a measure of temperature and humidity, are predicted to be some of the hottest globally due to relatively clear summer skies, low surface albedo of the Gulf, and a relatively shallow air boundary layer that concentrates heat and water vapor close to the surface (Pal & Eltahir, 2016). This translates to high summer water temperatures in the Gulf itself. Temperatures and salinities in the Gulf vary seasonally reaching extremes in the winter and summer. Temperatures can reach above 36° C and can fluctuate seasonally up to 20° C as it is relatively shallow and mostly enclosed (Burt *et al.*, 2011, Burt *et al.*, 2008, Coles & Riegl, 2013, Sheppard, 1993). With higher salinity and fluctuating temperature ranges above average of typical coral reefs (Coles & Riegl, 2013, Kinsman, 1964, Kleypas *et al.*, 1999), and relatively recent coral cover due to exposure due to glaciation events during the late Pleistocene epoch (Sheppard, 1993), corals have had to adapt to these relatively extreme conditions in a short period of time (D’Angelo *et al.*, 2015).

This study examines the effects of temperature tolerance on physiologies of corals in the Gulf. As *Symbiodinium* type can increase thermal tolerance (Hume *et al.*, 2013). This experiment utilized the aposymbiotic larvae created from controlled crosses of adults collected from a single population in the Gulf (Abu Dhabi, UAE (See for a detailed site description: Howells *et al.*, 2016)). This allows the effect of the host to be disentangled from any tolerance provided by the symbiotic complement. Here we estimated the heritability of thermal tolerance from larvae resulting from these crosses to determine the adaptive potential of this population using a quantitative genetics approach. The potential influence of parental genotype on larval thermal tolerance was also compared by regressing parental genotype and heat-exposed mortality using SNP markers throughout the genome. Finally, in an effort to understand possible causes of differential mortality in response to both temperature and salinity stressors, we compared transcriptional response in heat-tolerant and - susceptible families.

## Methods and Materials

### Experimental crosses and sampling

To study genetic variation in thermal tolerance of the uniquely heat-tolerant corals of the Gulf, we conducted a series of experimental crosses to produce larvae with known genetic relationships, following the same general procedures previously described for controlled crosses in corals (Howells *et al.*, 2016, Meyer *et al.*, 2009b). Briefly, we collected colonies of *Platygyra daedalea* prior to the annual spawning in April 2014 (Howells *et al.*, 2014) from a previously described population near Abu Dhabi, UAE (Howells *et al.*, 2016). We maintained colonies in recirculating aquaria containing 26° C, 42 ppt seawater to match conditions at the collection site. Nightly and prior to dusk, colonies were isolated into buckets to prevent cross-fertilization. When spawning occurred, we collected gamete bundles, separated egg and sperm bundles using a fine (50 μm) nylon screen, and prepared controlled crosses according to the design shown in Figure 1. We developed this crossing design to maximize the number of genetic crosses within the constraints of space and numbers of gametes available from each colony. After fertilization and rinsing away sperm, embryos were stocked into 2-l culture vessel containing freshly filtered seawater at a density of approximately 10 ml^−1^. We maintained larvae in culture for 7 days with complete water changes at 2-d intervals.

**Figure 1.**
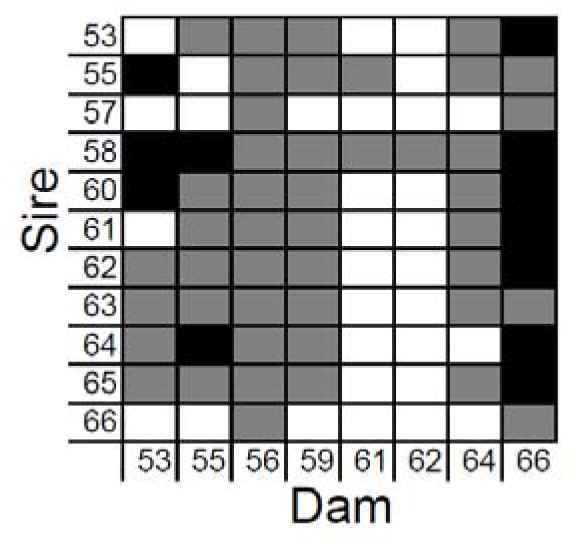
Design of experimental crosses used to produce coral larvae for quantitative genetic analysis of variation in thermal tolerance. Grey cells indicate successful crosses that produced sufficient larvae for experiments. Black cells indicate failed crosses, and white cells indicate crosses that were not attempted because of limited gamete numbers.

### Thermal stress experiments and measurement of thermal tolerance

To measure variation in thermal tolerance, we exposed larvae to elevated temperature stress treatments and documented their survival. For this analysis, we produced a series of 62 crosses using 11 sires and 8 dams (Figure 1), of which 57 families survived in sufficient numbers by 3 days postfertilization to be used in thermal stress experiments. We manually transferred 10-20 larvae from each family into individual wells of 6-well-plates, counting each group as we transferred them, with each well receiving on average 13.8 ± 4.5 larvae. We plated three replicate wells from each family, in each of two temperature treatments (26°C and 36°C), and took initial photographs of each well to determine initial sample sizes. One of the incubators was kept at 26° C and the other was ramped 2° C hr^−1^ to a maximum temperature of 36° C.

To quantify mortality resulting from these treatments we took multiple photographs of each well at several time points during this thermal stress experiment (15, 22, and 38 hours). Soft-bodied coral larval rapidly lyse and disintegrate after death, making direct counts of surviving larvae an effective procedure for measuring mortality in coral larvae (Dixon *et al.*, 2015). To confirm the accuracy of these photographs for counting larvae, we conducted counts by hand for 20 wells stocked with various numbers of larvae, and compared these with independent counts based on photographs of the same wells. These counts were highly correlated (R^2^= 0.96, df=18, p<1.0x10^−16^), so we based all subsequent analysis on counts from photographs.

We calculated mortality for each well and time point from the number alive in a well relative to the initial number in that well. To develop a metric for comparing the overall survival curves across families, we calculated Kaplan-Meier cumulative probabilities of survivorship for each well from the individual count data (Rich *et al.*, 2010).

### Analysis of genetic variation in thermal tolerance

The adaptive response to selection depends in part upon the narrow-sense heritability (*h^2^*) of the trait under selection, a measure of the relative contribution of genetic variation to variation in the trait (Falconer & Mackay, 1996). To understand the potential for adaptive responses to selection in already heat-adapted populations of *P. daedalea*, we estimated the heritability of thermal tolerance using a mixed model analysis of mortality data. For this analysis we explored both Bayesian and (Restricted Maximum Likelihood) REML approaches to estimate genetic variance components.

For the Bayesian analysis we analyzed variation in percent mortality data in each time point and temperature treatment separately, with random effects of animal and plate in the R (v3.3.1) package MCMCglmm (Hadfield, 2010). For these models we chose a weakly informative inverse Wishart prior (V=1, nu=0.002) (Wilson *et al.*, 2010) and conducted 10^7^ iterations after removing the first 10^6^ as burnin, then sampled every 100 iterations after ensuring this did not lead to autocorrelation. We conducted 10 replicate runs to ensure convergence on a common estimate. We also used the Bayesian approach to estimate heritability of Kaplan-Meier estimates of cumulative probability of survival in both the constant and elevated temperature treatments. We estimated heritability (*h^2^*) and 95% confidence intervals as described in Wilson et al. (2010).

For comparison with the Bayesian estimates of genetic variance components and heritability, we also analyzed the endpoint survival data (38 hr), fitting a similar animal model using a REML approach implemented in the software WOMBAT (version 27-06-2016). (Meyer, 2007). We calculated *h^2^* from these data as a ratio of additive genetic variance to total variance (Falconer & Mackay, 1996).

For comparison with these analyses of percent survival data and cumulative survival probabilities, we analyzed survival of individual larvae at the endpoint (38 hr) using a binomial regression with a logit link function similar in approach to previously published work for coral heritability estimates (Dixon *et al.*, 2015). In these models we included well and plate as random factors to account for shared effects of well and plate in the mixed model. As before, we chose a weakly informative prior with 10^7^ iterations, sampling every 100 iterations and removing the first 10^6^ as burnin.

Like most broadcast spawning corals (Baird *et al.*, 2009b) *P. daedalea* is a simultaneous hermaphrodite (Mangubhai & Harrison, 2008), so in experimental crosses using corals it is possible for a single colony to act as both a sire and a dam. However, some software for analysis of these crosses is unable to accept the same individual as both sire and dam, so the parents have to be re-coded in those programs depending on their use as sire or dam. To evaluate whether this software limitation affects conclusions drawn from these analyses, we analyzed percent survival data from the endpoint (38 h) in MCMCglmm coding parents both ways (the same name for both sire and dam, or different names for sire and dam) and compared the results.

### Testing for associations between paternal genotypes and offspring thermal tolerance

To investigate the genetic basis for the heritable variation in thermal tolerance identified in quantitative genetic studies, we tested for associations between parental genotypes and the thermal tolerance of their offspring. For this analysis, we conducted genome-wide SNP genotyping using 2bRAD, a sequencing-based approach (Wang *et al.*, 2012) that we’ve previously used for genetic analysis of corals (Dixon *et al.*, 2015) including *Platygyra daedalea* (Howells *et al.*, 2016). We analyzed the resulting sequences using a reference derived from aposymbiotic larvae (SRA accession SUB3015376) to exclude contamination from the algal symbiont (which we’ve previously shown to be present at negligible levels in these sequencing libraries anyway; (Howells *et al.*, 2016)). Finally, we tested for association between thermal tolerance of each larval family and their paternal genotypes at each locus, in an effort to identify loci and genomic regions associated with variation in corals’ thermal tolerance.

To determine parental SNP genotypes we extracted genomic DNA from each colony used as a parent in the crossing experiments. To minimize contamination from the algal symbionts, we lysed tissue samples with appropriate buffers for animal tissue and purified DNA from tissue lysates using the E.Z.N.A Tissue DNA extraction kit (Omega Bio-tek). To minimize enzymatic inhibitor that co-purify with genomic DNA from *P. daedalea* tissue (like many corals), we further purified the DNA by precipitating with isopropanol (final concentration 50%) at room temperature prior to preparation of sequencing libraries. We quantified DNA templates by fluorescence (Accublue; Biotium) and prepared sequencing libraries using 1 μg of DNA from each sample. We prepared sequencing libraries as previously described (Howells *et al.*, 2016, Wang *et al.*, 2012), using adaptors with fully-degenerate overhangs (5’-NN-3’) to capture all *Alf*I restriction fragments. We combined libraries in equimolar ratios based on qPCR quantification of the final sequencing libraries using primers ILL-Lib1 and ILL-Lib2 (AATGATACGGCGACCACCGA and CAAGCAGAAGACGGCATACGA respectively) for multiplex sequencing. We sequenced each library on the Illumina HiSeq 3000 platform (SR 50 bp reads) at Oregon State University’s Center for Genome Research and Biocomputing (CGRB), devoting approximately 1/30th of a sequencing lane (~10 million reads) to each sample.

Before calling genotypes from the resulting sequences, we processed raw sequences to exclude low-quality or uninformative reads. All bioinformatics analyses were conducted using the same pipelines we’ve previously described for 2bRAD genotyping (Howells *et al.*, 2016, Wang *et al.*, 2012), using scripts we’ve made publicly available at Github (https://github.com/Eli-Meyer/2brad_utilities/). We first truncated each read to exclude adaptor sequences, retaining the 36-bp *Alf*I restriction fragments. Then we excluded low quality reads (having more than 5 positions with quality scores <20) and reads matching oligonucleotides used in library preparation (Smith-Waterman alignment scores > 15). We analyzed sequence variation in these high-quality reads by comparison with a *de novo* reference produced using 2bRAD sequences prepared from aposymbiotic larval samples. We’ve previously developed these data for a genetic linkage map, and have archived the reads at NCBI’s Sequence Read Archive (SRA accession SUB3015376) to serve as a common reference across multiple studies. The *de novo* reference was produced as described (Wang *et al.*, 2012), by clustering high-quality reads from 2bRAD sequencing libraries to develop a set of consensus 2bRAD tags, each representing an *Alf*I restriction fragment from the *P. daedalea* genome, and is provided in Supplementary Information (Supplementary File S1).

To determine genotypes from these reads, we mapped HQ reads against this reference using SHRiMP (Rumble *et al.*, 2009), and filtered the resulting alignments to exclude weak (<32 matching bases) or ambiguous alignments (reads matching more than one tag equally well). Finally, we determined genotypes from the nucleotide frequencies observed at each locus, following the conservative frequency threshold approach we’ve previously described (Howells *et al.*, 2016, Wang *et al.*, 2012). Here we called loci homozygous if the frequency of a second allele was less than 1%, heterozygous if a second allele present at ≥25%, and left genotypes undetermined for loci with intermediate allele frequencies between those thresholds. After calling SNP genotypes for each sample, we compiled SNP genotypes from all samples into a single matrix, and filtered the combined genotypes to exclude loci sequenced at < 10× coverage in < 12 samples, and repetitive tags containing excessive SNPs (>2 SNPs per tag), and selected a single representative SNP from each of the tags that remained. The resulting set of high-confidence SNP genotypes was the basis for our analysis of associations between genotype and thermal tolerance.

To investigate the genetic basis for heritable variation in corals’ thermal tolerance, we used these SNP genotypes to test for associations between parental genotypes and the thermal tolerance of their offspring. We focused on paternal genotypes for this analysis, since maternal influences are likely to include non-genetic effects of egg quality. At each locus, we used a linear mixed model to test for associations between paternal genotypes and larval thermal tolerance, with dam included as a random factor to control for variation in egg quality. All statistical analyses were conducted in the R statistical environment, with linear mixed models in the lme4 package (Bates *et al.*, 2011). To correct for multiple testing, we controlled false discovery rate at 0.05 (Benjamini & Hochberg, 1995).

To investigate the biological consequences of these genetic variants, we compared the *Alf*I tag containing each SNP with *Alf*I tags identified from our reference transcriptome (above). We compared *Alf*I tag sequences from transcriptome and the 2bRAD de novo reference to identify the subset of tags that could be uniquely assigned to a single gene. For this analysis, we required at least 34 matching bases. We compared these overlapping tags (found in both the transcriptome and the 2bRAD reference) to identify the genes containing SNPs associated with thermal tolerance

To evaluate whether SNP genotypes from this analysis could be used to predict thermal tolerance of offspring based on paternal genotypes, we developed a simple model based on multilocus SNP genotypes. For each locus, we first identified the optimum paternal genotype (the genotype of sires producing the most heat-tolerant larvae). Then we classified each paternal colony based on the proportion of loci at which it harbored the optimum genotype. This produced a score ranging from 0-1, which we multiplied by the average mortality to predict relative mortality. To evaluate we the proportion of variation in mortality explained by these genotypes, we compared model predictions for each sire based on SNP genotypes with the average mortality observed in experimental crosses.

### Developing an annotated transcriptome database

To enable studies of gene expression in *P. daedalea*, we first developed an annotated transcriptome assembly to serve as a reference for expression analysis. For this purpose, we sequenced, assembled, and annotated cDNA libraries largely as described in (Kitchen *et al.*, 2015) with the following differences. We prepared cDNA using total RNA extracted from a pooled sample of approximately 20 aposymbiotic larvae from a single cross (55x58), and another pooled sample of approximately 20 eggs from the same dam (58). We extracted RNA from each sample using the Omega Bio-tek E.Z.N.A. RNA kit, followed by precipitation with LiCl (4 M final concentration). First strand synthesis was accomplished using Superscript II Reverse Transcriptase (Invitrogen Fisher Thermo), relying on template switching activity to add the custom RNA oligos to the 5’ end, then amplified and purified as described in Kitchen et al. (2015). To minimize the sequencing coverage wasted on highly expressed genes, we normalized the amplified cDNA as previously described for coral transcriptomes on an older sequencing platform (454) (Meyer *et al.*, 2009a), then re-amplified the normalized cDNA, and conducted the remainder of library preparation as described in (Kitchen *et al.*, 2015).

We sequenced this library in a single flow cell of paired-end 75-bp reads on Illumina MiSeq at Oregon State University’s Center for Genome Research and Biocomputing (CGRB). Prior to assembly, we filtered the raw reads for uninformative and low-quality reads as we have previously described (Meyer *et al.*, 2009a) using custom Perl scripts available on GitHub (https://github.com/Eli-Meyer/transcriptome_utilities). We then assembled the high-quality reads using Trinity v2.0.2 (Grabherr *et al.*, 2011) and annotated the resulting assembly with gene names and GO terms (Ashburner *et al.*, 2000) based on BLAST+ (Altschul *et al.*, 1990) comparisons with the UniProt database (version 2014_09). Finally, to optimize the accessibility of this resource by the coral research community, we developed a searchable database using SQLITE (https://www.sqlite.org/) and a simple PHP interface for searching this database, and hosted this resource on the author’s laboratory website at Oregon State University (http://ib.oregonstate.edu/~meyere/DB/Pdae/search.php).

### Profiling transcriptional responses to thermal stress

To further investigate functional differences between these phenotypes, we surveyed transcriptional responses to thermal stress in heat-tolerant and susceptible corals. For these experiments we used a subset of families, chosen based on availability of sufficient larvae to set up the experiment and sufficient yield in RNA extractions. To fit the experiment within budget constraints, we included progeny from 5 sires incubated at 26 and 36°C for a total of 13 samples.

This design allows us to estimate effects of sire as a main effect and its interaction with temperature, but not to make specific pairwise comparisons between families. We distributed approximately10-30 larvae (average = 16.9 ± 5.2) larvae from each family into 6-well-plates, and distributed the plates between two incubators. All larvae used for analysis of gene expression were incubated in 42 ppt seawater, the same salinity as the main cultures. (Additional larvae from each family were incubated in parallel using 35 and 46 ppt seawater for another experiment, and some of those samples were used below for analysis of allele frequencies). One incubator was held at ambient temperature (26° C) while the other was increased 2° C hr^−1^ to a maximum of 36° C. We photographed each well prior to temperature treatments and again at the conclusion of the experiment (24 hr). We sampled all animals at this time from both stress and control treatments and preserved them in RNAlater for gene expression analysis.

We extracted RNA from each sample (pools of ~10-20 larvae), further purifying these samples with LiCl to remove residual genomic DNA and enzymatic inhibitors that copurify with nucleic acids in these samples. Using these purified RNA samples (approximately 150-300 ng RNA per sample) we prepared tag-based cDNA libraries for gene expression as previously described for gene expression on Illumina HiSeq (Crowder *et al.*, 2017, Meyer *et al.*, 2011). To avoid distortion of transcript abundances in these libraries, we carefully minimized the number of PCR cycles during amplification, amplifying all samples for only 19 cycles. We added sample specific barcodes via PCR and size selected each library (retaining 250-350 bp fragments) using gel electrophoresis. Finally we combined libraries in equimolar amounts and sequenced the combined pool twice on single-end 75-bp reads (HiSeq 2500), at two different sequencing facilities (University of Texas - Austin, University of Chicago).

Prior to statistical analysis of gene expression, we first applied a minimum coverage threshold of ≥ 5 reads per sample to avoid low abundance transcripts where sampling error would obscure differential expression. Then we used a negative binomial model implemented in the DESeq2 package (Love *et al.*, 2014) to evaluate the effects of sire, temperature, and their interaction. We tested the significance of each effect by comparing full and reduced models using likelihood ratio tests (Marioni *et al.*, 2008). Dam was included in all models to account for effects of egg quality and maternal genotypes, handling this as a fixed factor because of the small number of dams used per sire. After differential expression analysis of each gene, we controlled for multiple tests to control false discovery rates at < 0.05 (Benjamini & Hochberg, 1995, Love *et al.*, 2014). All statistical analyses were conducted in the R statistical environment (R Core Team, 2013). Statistical tests were based on raw counts, but to visualize patterns of gene expression we plotted rlog transformations of the gene counts (Love *et al.*, 2014).

To evaluate the functional context of these gene expression patterns, we conducted a functional enrichment analysis based on Gene Ontology (GO) annotation of our reference transcriptome. We used the ErmineJ software (ver .3.0.2) (Gillis *et al.*, 2010, Lee *et al.*, 2005) to investigate enrichment of specific biological process terms in the genes differentially expressed in response to temperature and genotype as main effects and their interaction. We analyzed raw p-values from DESeq2 using receiver operator characteristics (ROC) (Gillis *et al.*, 2010). Each GO category was scored using Wilcoxon rank sum of all member genes normalized and fit to the ROC curves calculated to determine significance (Breslin *et al.*, 2004) and only those passing BH adjusted p-values (FDR<0.05) are reported. These tests provided an objective procedure for summarizing the functional interpretation of differential gene expression in our experiments.

In addition to information on expression levels, RNASeq data also provide information on sequence variation. To study relationships between thermal tolerance and sequence variation in expressed sequences, we re-analyzed alignments of cDNA from each sample against the annotated reference transcriptome. For this analysis we used all samples incubated at control temperatures (26°C) (including additional samples cultured at 35 or 46 ppt salinity as part of another experiment), to avoid distortions in allele frequencies that may result from thermal-stress induced mortality (24 samples altogether). To minimize false positives arising from highly expressed genes, we randomly resampled sequence alignments from each biological sample (SAM files) to set the maximum coverage at 100 reads per gene. Because our tag-seq libraries focuses sequencing effort on a narrow window of each transcript (Meyer *et al.*, 2011) this threshold (number of reads) is approximately equal to the total alignment depth. We processed alignments using samtools and bcftools (http://www.htslib.org) to sort and index alignments, and compile information on allele frequency at each locus in VCF format. We compiled these data from each sample into an allele frequencies matrix and filtered the dataset prior to statistical analysis to minimize missing data and focus on informative loci. We excluded low-coverage loci genotyped in too few samples (requiring > 20× coverage in > 4 samples). The filtered set of loci resulting from this process were used for statistical analysis of allele frequencies. Since each sample consists of pooled RNA from multiple full-sibling larvae, we analyzed allele frequencies in each sample directly rather than attempting to call genotypes from pooled samples. We considered a locus polymorphic if a second allele was detected in at least 5 reads for at least 5% of samples.

To investigate relationships between thermal tolerance of each family and allele frequencies at each SNP locus, we used logistic regression (glm: logit link function) of the counts of each allele in each sample on the family-specific measurements of thermal tolerance (mortality at elevated temperature). Statistical tests were conducted in R. We corrected for multiple tests by controlling false discovery rate at 0.05.

## Results

### Genetically determined variation in corals’ thermal tolerance

Larval mortality during thermal stress treatments varied substantially as a function of heat, genotype, and their interaction. Larvae that were in the heated treatment were 30.6 times more likely to die than in those in the control temperature (*P* < 10^−16^). In the control conditions, average mortality remained low overall, averaging 11.1% (±13.2%) across all 57 families at the end point (38 hr). In contrast, mortality in the heat treatment increased substantially over time, averaging only 6.4% (±18.9%) at 15 hr, but reaching 34.9 (±23.8%) and 64.5% (±29.8%) at 22 and 38 hrs, respectively. When controlling for dam, there was also a significant effect of sire. In contrast to the relatively constant mortality at control conditions, mortality varied widely among sire at the elevated temperature (Figure 2). Compared to highest performing sire (65: avg. mortality=10.1±12.3%), offspring sired by colonies 61 and 64 were 13.5 and 17.4 times likely to die in the elevated temperature treatment, respectively (Figure 2).

**Figure 2:**
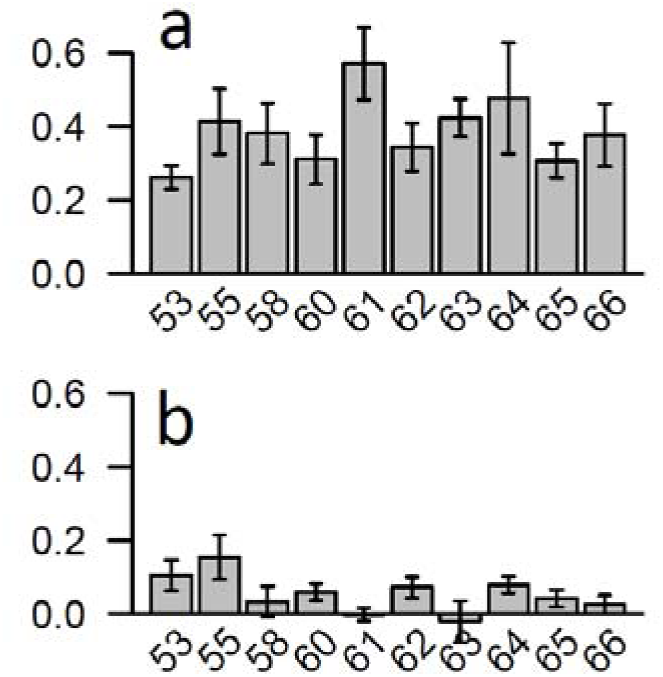
Coral larvae from experimental crosses vary widely in thermal tolerance. Each bar represents the average mortality of all crosses from a single sire, and error bars show standard error of the mean. a) Mortality in larvae incubated at 36°C for 38 hours. b) Mortality in control incubations of the same families at 26°C.

To quantify the heritability of this variation in thermal tolerance, we partitioned variation in mortality using an animal model. We first estimated heritability of thermal tolerance using direct counts of the number of animals surviving in each well at the experimental endpoint (38 hr) in a Bayesian logistic regression model (MCMCglmm). This analysis revealed that in control conditions, genetic variation explains very little of the variation in mortality (*h^2^* = 0.0007; Table 1). In contrast, variation in mortality during thermal stress was much more heritable (*h^2^* = 0.484; Table 1). This value of *h^2^* suggests substantial potential for adaptive responses to selection for increased thermal tolerance. There was high precision in the estimates when the model was rerun indicating convergence on a stable heritability.

**Table 1.**
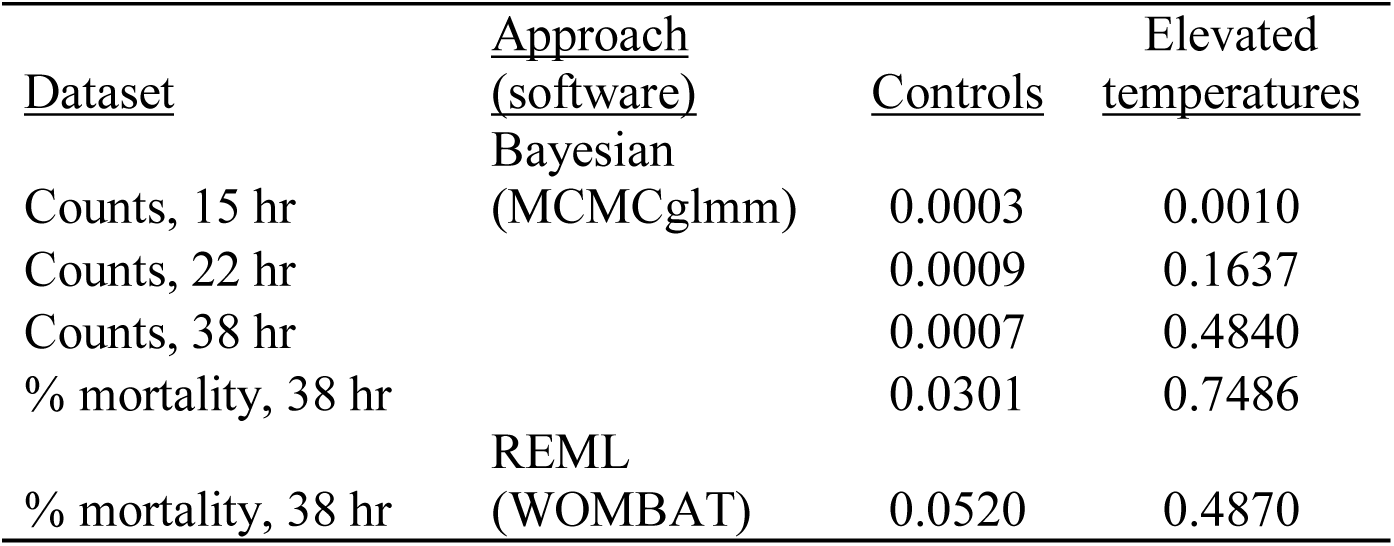
Estimates of narrow-sense heritability variation in corals’ thermal tolerance based on mortality in control (26°C) and elevated temperatures (36°C).

To evaluate whether this conclusion is robust to decisions about measurement or analysis of mortality data, we compared *h^2^* estimates from several different analyses, which consistently showed that variation in mortality during thermal stress was highly heritable in these crosses (Table 1). Sampling time influenced *h^2^* estimates, since mortality increased over time. While heritability of mortality remained low (0.0003 - 0.0009) throughout the experiment in the control conditions, where mortality was low and relatively constant, *h^2^* of mortality increased over time (from 0.001 to 0.48) in the thermal stress treatment as overall mortality increased. To develop an aggregate measure of mortality incorporating all timepoints, we estimated Kaplan-Meier survivorship for each family, and *h^2^* values estimated from this trait were comparable to the endpoint (38 hr) analysis of counts: low in control temperatures (*h^2^*=0.048; Table 1) and high in thermal stress treatments (*h^2^*=0.755; Table 1).

We found that these conclusions were similarly robust to decisions made during analysis, including the modeling approach (REML versus Bayesian inference) and data type (mortality counts versus percent mortality). Analysis of percent mortality in each well produced higher estimates of *h^2^* than the logistic regression models based on counts of larval mortality (above), but supported the same general conclusions. This approach yielded low estimates of *h^2^* for mortality in control conditions (0.03) and high *h^2^* for mortality in thermal stress (0.749). Analysis of the endpoint mortality data (as percent mortality) using a REML model in the software WOMBAT produced slightly different estimates but again supported the same general conclusions. These models revealed low and non-significant heritability in control conditions (*h^2^* = 0.052 ± 0.128; *P* = 1) but moderate and highly significant heritability in thermal stress (*h^2^* = 0.487 ± 0.210; *P* = 0.00007). These results were also robust to decisions made within this analysis. The decision of whether to code corals as the same individual or different when using the coral as a sire or a dam had little effect on *h^2^* estimates (*h^2^* = 0.749 when coded the same, *h^2^* = 0.707 when coded differently). Overall, our analysis robustly supports the conclusion that the populations sampled to produce these crosses harbor substantial natural variation in thermal tolerance, which in principle might support adaptive responses to rising ocean temperatures.

### Associations between paternal genotypes and thermal tolerance of offspring

To further investigate the genetic basis for these heritable differences in thermal tolerance, we tested for relationships between multilocus SNP genotypes and thermal tolerance in each family. To that end we genotyped the parents from these crosses with a sequencing based approach (2bRAD), recovering 4.7 million reads on average per sample, most of which (95.7% on average) passed quality filtering and adaptor screening. All sequencing yield and quality statistics are reported in Supplementary Table S2. On average, we genotyped 1.38 Mb in each of the 24 parental colonies. After combining these SNP genotypes and filtering the genotype matrix to minimize missing data and genotyping errors, we recovered 20,507 high-quality SNPs for genetic analysis.

We used these SNPs to estimate genetic distances among individuals (one minus the proportion of shared alleles), revealing genetic distances among colonies ranging from 0.129 to 0.209 and confirming that no colonies used in this study were identical genotypes (which can occur in some corals through fragmentation or other asexual processes). Genomewide estimates of individual heterozygosity in each sample ranged from 0.0013 to 0.0024. On average, 23% of loci that were polymorphic within the population were heterozygous in each individual.

To identify markers and genomic regions associated with the genetic variation in thermal tolerance documented in this study, we used these parental genotypes to test for relationships between thermal tolerance and genotype at each locus. Because our crossing design provided reasonable replication of sire effects across multiple dams, and because paternal contributions are largely genetic while maternal contributions often include confounding effects (e.g. energetic content of eggs, or mitochondrial traits), we focused on sire effects for this analysis. We used linear mixed models to test for associations between thermal tolerance and paternal genotype at each locus, including random effects of dam to account for variation in egg quality or maternal genotypes. We tested these genotype-phenotype relationships at 11,925 loci and identified 131 SNPs significantly associated with thermal tolerance after multiple test correction. Effect sizes of these loci were generally small, with each SNP explaining on average 12% of variation in thermal tolerance among families (marginal R^2^; range = 8 to 18%). To illustrate the scale of these effects, several examples of markers significantly associated with thermal tolerance are shown in Figure 3a,b. The complete list of 2bRAD markers significantly associated with thermal tolerance is shown in Supplementary Table S3.

**Figure 3:**
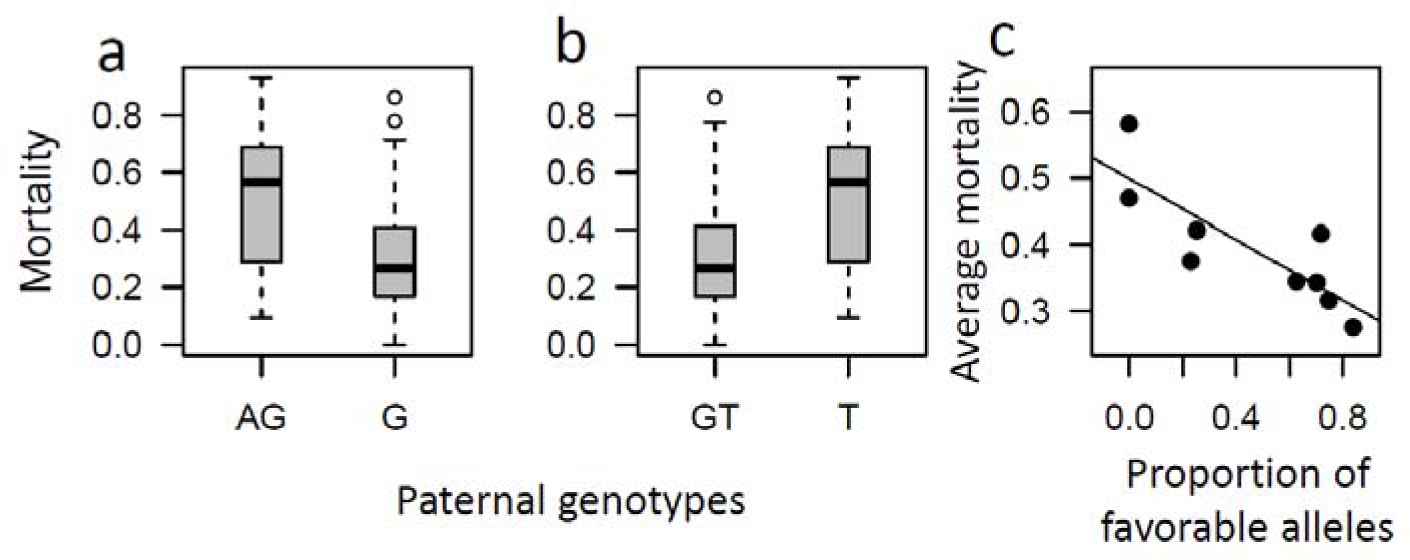
Our analysis identified genetic markers where paternal genotypes are associated with thermal tolerance of their offspring. The first two panels show arbitrary examples of individual loci showing significant relationships with thermal tolerance in a mixed model analysis. Altogether we identified 131 markers significantly associated with mortality. a) denovoLocus21161, position 11. b) denovoLocus4455, position 31. Panel (c) shows the results of a multilocus model using SNP genotypes to predict the average thermal tolerance of each sires’ offspring from paternal genotypes at all markers identified here. This panel of SNP genotypes explains 69% of variation in mortality during thermal stress (P = 0.0056, R2 = 0.689).

These genetic markers provide a potential tool for prediction of corals’ thermal tolerance from SNP genotypes of parental colonies. To evaluate the overall contribution of paternal genotypes to thermal tolerance of their offspring, we calculated the proportion of loci at which each father harbored the optimum genotype (based on comparisons across all families at each locus). We found that paternal genotype explains 68.9% of variation in thermal tolerance of their offspring, across all 131 significant SNP markers (Figure 3c). We also estimated genetic distances among parents to evaluate whether thermal tolerance is associated with genomewide heterozygosity, but found no relationship (R^2^=0.001, p=0.84).

### Assembly and annotation of a reference transcriptome for gene expression analysis

The annotated transcriptome we produced using normalized cDNA from aposymbiotic larvae and eggs includes 51,200 contigs. Like most *de novo* transcript assemblies this includes a substantial number of small contigs that skew the overall distribution: 48.8% of contigs are <400 bp, so the average length and N50 of the initial assembly are relatively low (542 and 648 respectively). Excluding these short sequences improves the length distributions (average=793, N50=862), emphasizing that the assembly includes a reasonable number of long transcripts (26,195 > 400 bp, 15,927 > 600 bp, and 9,891 > 800 bp). The initial assembly is also somewhat redundant, with clusters of related transcripts recorded as components by the assembler. The 51,200 contigs include 42,701 components (21,380 > 400 bp, 12,985 > 600 bp, and 8,121 > 800 bp).

Functional annotation based on BLASTX comparisons with the UniProt database identified matches for relatively few of these (14,646 transcripts). This primarily reflects the gene content of long transcripts, since short sequences are less likely to include regions of local similarity (40.8% of transcripts ≥ 400 bp have a significant BLAST match, compared with 15.7% of transcripts < 400 bp). For a subset of these (10,375 transcripts), we were able to assign GO annotation to the transcripts based on GO annotation of their best BLAST match. This provided a resource for functional enrichment analysis of gene expression profiles using this reference.

To evaluate the gene content and completeness of the transcriptome we conducted several sequence comparisons. First we compared the transcripts with a collection of 458 orthologous groups of proteins widely conserved among eukaryotes, the CEGMA database (Parra *et al.*, 2009). Our transcriptome assembly includes significant matches (bitscores > 50) for 442 of those (96.5%), suggesting that the majority of conserved eukaryotic genes are captured in our assembly. Comparison with gene models from the sequenced coral *Acropora digitifera* tempered this conclusion, showing that while many transcripts (18,199) matched predicted proteins from the coral genome, these included matches for only 10,044 of the 23,677 *A. digitifera* proteins (42.4% of the total). Based on the length distribution of our assembly, these sequence comparisons, and the expectation that only a subset of genes are expressed in the developmental stages sampled here, we conclude that this transcriptome assembly probably includes approximately half of the ~24,000 genes expected in a coral genome.

Finally, to evaluate the completeness of our transcript assemblies we compared protein translations from each transcript with the lengths of their best match in the A. digitifera genome, to estimate ortholog hit ratios (OHR), a metric describing the proportion of each gene represented in the corresponding transcript (O’Neil & Emrich, 2013). Of the 10,044 coral genes (*A. digitifera*) matched by transcripts in our assembly, the assembly included one or more reasonably complete transcripts for 3,847 of these. Of course, as the length distribution suggests, the assembly also includes only partial transcripts for many genes (for 6,196 genes, the maximum OHR in our transcripts was <0.7). Although incomplete (like all or most *de novo* reference transcriptomes), this annotated transcriptome assembly provides a useful reference for gene expression analysis and other applications. To maximize the accessibility of this resource for the coral research community, we’ve provided the database in a searchable form on the author’s website at (http://ib.oregonstate.edu/~meyere/DB/Pdae/search.php).

### Analysis of gene expression in heat-tolerant corals

For additional insights into the functional basis for differences between heat-tolerant and heat-susceptible we compared gene expression profiles in these phenotypes. We profiled gene expression in larvae from five different sires, testing for effects of sire, temperature, and their interaction in a negative binomial model. Here we describe an overview of these expression profiles; the complete list of differentially expressed genes is shown in Supplementary Table S4.

We found that 196 genes were differentially expressed in thermal stress treatments, regardless of parentage (FDR < 0.05) (Figure 4). These included two major clusters of expression patterns indicated in Figure 4a. The 130 genes in cluster 1 were up-regulated during thermal stress in all families, and this list includes genes with known roles in transcriptional responses of corals to thermal stress (heat shock proteins 19.8, 70, 90, and 90a1; four calumenin genes, two endoplasmin genes, and a fluorescent protein). The 66 genes in cluster 2 were down-regulated during thermal stress. Many of these genes have also been previously described in transcriptional stress responses of corals, including ferredoxin, peroxiredoxin, peptidyl-prolyl cis-trans isomerase, calmodulin, and glutathione synthetase. Enrichment analysis of temperature effects identified two biological functions that were significantly enriched in responses to elevated temperatures: protein folding (GO:0006457; adjusted p-value = 0.039) and response to stress (GO:0006950; adjusted p-value = 0.0025), providing objective support for the processes highlighted here. The complete list of significant results from functional enrichment analysis is shown in Supplementary Table S5.

**Figure 4:**
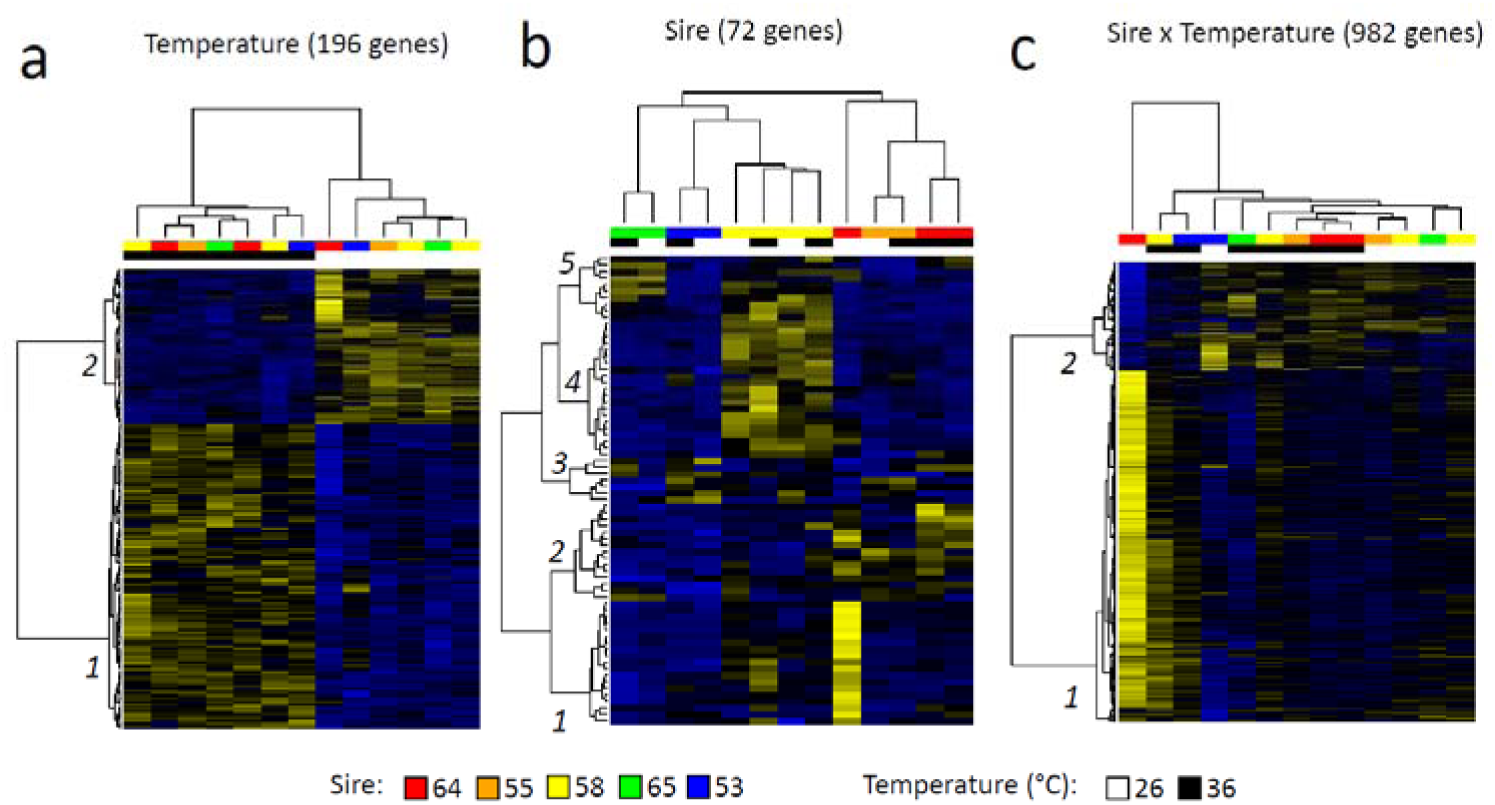
Corals’ transcriptional responses to thermal stress are strongly influenced by parentage. These heatmaps show genes that were differentially expressed as a function of (a) temperature treatment, (b) sire, and (c) their interaction. The sire (paternal parent) and temperature treatments are color coded in a bar above each heatmap.

In contrast to the robust transcriptional responses induced by thermal stress, fewer genes (n=72) were constitutively expressed at different levels as a function of sire (FDR < 0.05) (Fig 4b). Although fewer in number, these expression profiles showed clear signals of sire on gene expression, with progeny of each sire expressing a cluster of genes more highly than progeny of other sires. Interestingly, the two families with highest thermal tolerance (progeny of sires 53 and 58, blue and yellow in Fig 4b, with < 20% mortality each) clustered together in this analysis, as did the two families with the lowest thermal tolerance (progeny of sires 55 and 64, orange and red in Fig 4b, with > 70% mortality each). Like the transcriptome overall, a large fraction of these DEG were not annotated (60/72 lacked annotation), and the functional interpretations of the few annotated genes in each cluster are unclear. Despite these limitations, there were a number of GO terms enriched in both the biological and molecular function categories (Supplemental Table S5). These included DNA metabolism and synthesis pathways (GO:0006259, GO:0071897, GO:0034061), translation (GO:0006412, GO:0043043) and reverse transcription (GO:0006278: see below). These comparisons demonstrate substantial differences in constitutive levels of gene expression between progeny of different sires, highlighting a mechanism that may contribute to the differences in thermal tolerance among these families.

Most of the differential expression we found in these data (982 DEGs) were temperature by sire (TxS) interactions, indicating genes where the transcriptional response to elevated temperatures differed between families (Fig 4c). Enrichment analysis of the sire × temperature interaction identified several processes enriched in these DEGs, including DNA metabolism (GO:0006259; adjusted p-value = 0.000039) and receptor-mediated endocytosis (GO:0006898; adjusted p-value 0.0089).

We identified two clusters of expression patterns in these interactions, designated 1 and 2 in Fig 4c. The 751 genes in cluster 1 were expressed at relatively low levels in most samples independent of temperature, while the most heat-tolerant families (progeny of sires 53 & 58) up-regulated these genes during thermal stress, and the most heat-susceptible families (progeny of sire 64) down-regulated these genes. This gene list was dominated by a number of functional categories not typically associated with thermal stress responses, with functional roles that suggest differences in DNA replication. Three of these DEGs matched histones. Twenty-three DEGs matched reverse transcriptase sequences from other marine invertebrates that may represent viral or transposable elements. Consistent with this idea, cluster 1 also included 7 DEGs annotated as integrase or recombinase, and 7 annotated as transposase.

The 231 genes in cluster 2 showed the opposite pattern, being expressed at relatively high levels in most samples independent of temperature, while a few crosses up-(sire 64) or down-regulated (sires 53 & 58) these genes during thermal stress. This cluster included several genes previously reported in association with thermal stress responses of corals. These include thioredoxin, hemicentin, fluorescent protein, tachylectin, and galaxin. Another notable pattern in cluster 2 was the abundance of genes associated with ribosome synthesis (8 DEGs matching ribosomal proteins or RP kinase).

While the number of families in this analysis is inadequate to draw general conclusions about relationships between expression levels and thermal tolerance, our finding of extensive sire x temperature effects on gene expression in families with contrasting thermal tolerances suggests that variation in transcriptional stress responses contributes to variation in corals’ thermal tolerance phenotypes. The genes identified here suggest that variation in corals’ thermal tolerance phenotypes may result in part from variation in thermal stress responses, capacity for protein synthesis, and the activity of viral or transposable elements.

### Analysis of transcript sequences in heat-tolerant corals

The same RNASeq data used for gene expression analysis also provided information on sequence variation among families. After estimating allele frequencies from RNASeq alignments in all samples from the control temperature (n=24), combining these into an allele frequencies matrix, and filtering for coverage (≥20×) and missing data (<10%), we were able to estimate allele frequencies at 220,912 basepairs distributed across 5,892 different transcripts. The final dataset included 7,004 SNPs distributed across 2,407 transcripts.

To evaluate associations between genetic variation in expressed sequences and thermal tolerance of each family, we conducted linear mixed model analysis at each SNP locus. We focused on larvae held at control temperatures for this comparison to avoid distortions in allele frequencies that may result from thermal-stress induced mortality. We tested for associations using logistic regressions, and found that 586 SNPs showed significant associations between allele frequencies and mortality during thermal stress. Examples of these relationships are shown in Figure 5, and the complete list of sequence variation significantly associated with thermal tolerance is shown in Supplementary Table S6.

**Figure 5:**
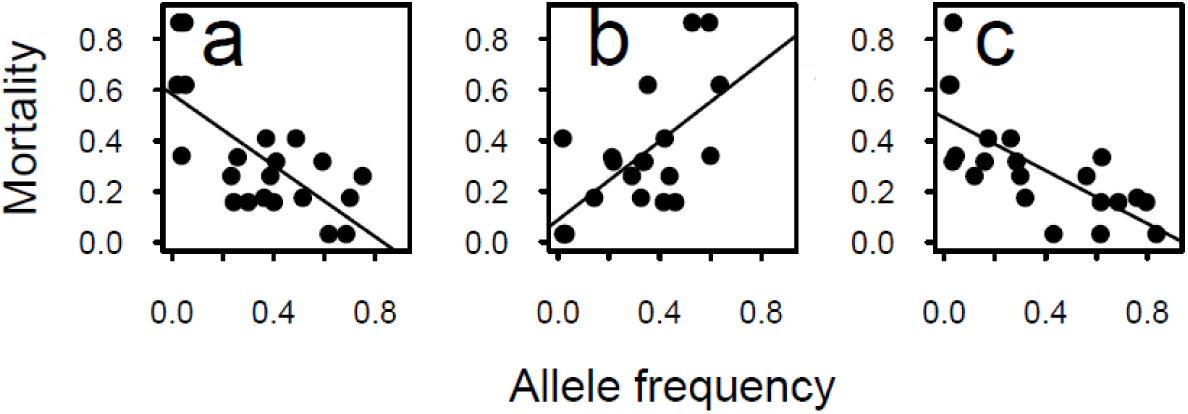
Genetic variation in expressed sequences is associated with variation in thermal tolerance. The figure shows three example transcripts out of the 586 markers where allele frequencies identified from RNA-Seq analysis were significantly associated with thermal tolerance (logistic regression; FDR < 0.05). a) c8557_g1_i1, a transcript matching Uniprot accession C3ZWH9 (an iron-sulfur domain containing protein). b) c11375_g1_i1, a transcript matching T2M547 (Regulator of nonsense transcripts). c) c5214_g1_i1, a novel transcript lacking matches in Uniprot.

The significant SNPs were distributed among 421 different transcripts. The majority of these (66%) lacked functional annotation, but we further investigated the functional annotation of the 144 annotated transcripts to investigate which categories of genes have sequence variation associated with thermal tolerance in corals. Overall, these genes were distributed across several core aspects of cellular metabolism. One core category was the regulation of transcription and RNA processing. The list includes multiple genes associated directly with transcription (e.g. Mediator of RNA polymerase II transcription subunit 11, RNA polymerase II-associated protein 3), with RNA processing (e.g. rRNA-processing protein EFG1, U3 small nucleolar RNA-associated protein 6-like protein, Pre-rRNA-processing protein TSR1 like protein, U6 snRNA-associated Sm-like protein LSm6, mRNA cap guanine-N7 methyltransferase), or tRNA synthesis (Serine—tRNA ligase, Asparaginyl-tRNA synthetase). There are also several transcription factors or regulators of transcription factors (e.g. Heat shock factor binding protein 1-like protein, Hes4, Out at first protein, and Forkhead box protein K1)

Another dominant category was protein metabolism. The list includes genes with roles in protein synthesis (e.g. ribosomal proteins L4, L15, S14, S17, S27, and P1), and protein degradation (ubiquitin, two ubiquitin-protein ligase genes, Cathepsin L protease, CAAX prenyl protease, and two proteasome subunit genes). Finally, we also found genes in processes previously implicated in thermal stress responses of corals. These included oxidative stress responses and cellular redox homeostasis. The list includes glutaredoxin, superoxide dismutase, and two glutathione S-transferase genes. Other genes were associated with unfolded protein responses, including chaperone proteins Hsp90 and universal stress protein.

In species with sequenced genomes, genomic association studies often aim to identify a region of the genome (and a set of genes) associated with variation in the trait of interest. The absence of a sequenced genome for *Platygyra daedalea* limits the functional inferences that can be drawn from these data, but 2bRAD tags are randomly distributed across the genome so that some match genes directly. To identify these genes containing SNPs associated with thermal tolerance, we first extracted *Alf*I tags from the annotated transcriptome assembly (described above) and aligned tags from our 2bRAD reference against the transcriptome tags, requiring ≥34 matching bases out of 36. This procedure identified 4,909 2bRAD tags associated with contained within genes. This subset of tags included 1,049 SNPs, and 5 of these were significantly associated with thermal tolerance. While this sampling of genes is obviously too incomplete for any general conclusions, the genes identified in this analysis make sense from the perspective of coral thermal tolerance. Two are involved in regulation of transcription (RNA Polymerase and Forkhead, a transcription factor), suggesting that genetically-determined variation in transcriptional responses may play a role in the observed variation in thermal tolerance. Another gene identified here (UFD1) is associated with ubiquitin-dependent protein degradation, suggesting that genetically-determined variation in this process may play a role in variation in corals’ thermal tolerance. The remaining two transcripts lacked functional annotation.

Together, these findings from genetic analysis of the parents are consistent with the pattern emerging from analysis of gene expression. Genetic variation in corals’ thermal tolerance is associated with variation in transcriptional responses that may be driven in part by variation in these regulatory genes.

## Discussion

Our study has demonstrated that coral populations in the warm waters of the Gulf harbor extensive genetic variation in thermal tolerance. We identified genetic markers for thermal tolerance in these populations, and found that these markers explain a large fraction of the observed variation in thermal tolerance. Transcriptional responses to thermal stress are highly influenced by genetics, and these effects include core cellular processes of DNA and protein metabolism in addition to known components of coral stress responses. We found that sequence variation in transcripts was also associated with thermal tolerance in these crosses, implicating a similar list of cellular processes as differential expression analysis. These findings suggest several interesting implications for coral adaptation and the functional basis of variation in corals’ thermal tolerance.

Our study of corals in the world’s warmest reefs found that a large fraction of variation in thermal tolerance was attributable to genetic variation (narrow-sense heritability, *h^2^*). Estimates of heritability of thermal tolerance after 38 hrs of exposure at high temperatures range from moderate to high (0.48-0.7) depending on the method utilized. This suggests substantial potential for adaptive responses to selection for thermal tolerance in these populations. Importantly, heritability is a function of genetic and environmental variation, and cannot be generalized across populations (Visscher *et al.*, 2008). Documenting high heritability of thermal tolerance of corals adds an important data point to a spare literature that currently includes only Acroporids (Dixon *et al.*, 2015, Meyer *et al.*, 2009b, Palumbi *et al.*, 2014). Here we describe comparable levels of heritable variation in thermal tolerance in a distantly related Robust coral, suggesting comparable potential for adaptive responses.

As corals face warming oceanic conditions, it may be possible for populations already experiencing high temperatures to supply heat-tolerant larvae to other regions (e.g. Kleypas *et al.*, 2016). As the dispersal stage of scleractinian corals, planula larvae help by bringing in novel genotypes that could confer higher thermotolerance. This assumes that 1) larvae will immigrate to reefs in need of larvae, 2) larvae from a given population will have similar thermotolerance to their parents, 3) reefs with warmer water temperatures will spawn larvae that also confer these benefits, and 4) recruits will survive in these new environments. In a recent simulation study, source reefs in the coral triangle were able to transport larvae to reefs that had different thermal regimes in the process introducing novel alleles conferring beneficial survival (Kleypas *et al.*, 2016). Although some Scleractinian corals have high larval longevities (Graham *et al.*, 2008, Wilson & Harrison, 1998), *P. daedalea* is competent to settle after just 2.5 days with important ramifications for determining pelagic larval duration (Miller & Mundy, 2003). Although there is some evidence based on a single microsatellite marker that *P. daedalea* populations are structured in Australia (Miller & Ayre, 2008), there is evidence of connectivity between reefs at a scale of 10s of km in a congener (*Platygyra sinensis*) near Singapore (Tay *et al.*, 2015). Thus it may be possible for the thermally tolerant corals in the Gulf to supply tolerance alleles to other populations.

The observation of highly heritable variation in thermal tolerance in Gulf corals also raises an interesting question. Despite strong selection for thermal tolerance, extensive genetic variation in thermal tolerance remains in these populations. Adaptive responses to selection are expected to deplete genetic variation in the trait under selection (Falconer & Mackay, 1996), but clearly this has not occurred in these populations. This may result from weak selection for thermal tolerance in nature, but empirical comparisons of larval thermal tolerance and environmental conditions suggest selection for thermal tolerance is strong in these populations (Howells *et al.*, 2016).

Gene flow from outside the Gulf may contribute to this; the relatively low differentiation we have previously documented suggests ongoing gene flow between these populations (Howells *et al.*, 2016). This gene flow may act to limit local adaptation in the Gulf. Balancing selection may also play a role. This may result from the extreme seasonal variation in temperatures in this region (Howells *et al.*, 2016). Coral populations inhabiting variable environments have been shown to be more tolerant than populations that experience more consistent conditions (Barshis *et al.*, 2013, Kenkel & Matz, 2016, Oliver & Palumbi, 2011, Schoepf *et al.*, 2015). Tradeoffs between maximum and critical temperature tolerance are widespread in animal physiology, suggesting that selection for high critical maximum temperatures during summer may be balanced by selection for low critical minimum temperatures. Other unknown correlations among traits may act in the same way, limiting adaptive responses to selection for high thermal tolerance and maintaining genetic variation in this trait.

Multilocus SNP genotyping explained a large fraction of thermal tolerance in our experimental crosses. If these markers are transferable across populations, the relationship shown in Figure 3c may allow prediction of genetic variation in thermal tolerance based on SNP genotyping. This would be enormously valuable for conservation and restoration efforts, since sampling and SNP genotyping are orders of magnitude less difficult and costly than empirically measuring thermal tolerance in larval stages. Such markers would be directly applicable for assisted translocation efforts aiming to provide heat-tolerance alleles to populations lacking those alleles (Coles & Riegl, 2013). Transferable genetic markers for thermal tolerance would allow restoration efforts to enrich thermal tolerance alleles in their outplanted population or parental colonies used for propagation. Recently, it has been suggested that assisted evolution may be required for maintenance of coral cover on reefs whether by transport of adult colonies or spawned larvae with or without settlement assistance (van Oppen *et al.*, 2017). The estimates of quantitative genetic parameters described here, and the specific genetic markers we’ve developed, will be important tools for efforts in this direction. Future studies should evaluate the generalizability of these markers across populations to evaluate the feasibility of applying these markers in conservation and restoration efforts.

Our studies also provide insights into the functional basis for variation in corals’ thermal tolerance. While the number of families in this analysis is inadequate to draw general conclusions about relationships between expression levels and thermal tolerance, our finding of extensive sire × temperature effects on gene expression in families with contrasting thermal tolerances suggests that variation in transcriptional stress responses contributes to variation in corals’ thermal tolerance phenotypes. The genes identified here also suggest that variation in corals’ thermal tolerance phenotypes may result in part from variation in thermal stress responses, capacity for protein synthesis, and the activity of viral or transposable elements. We found extensive variation in gene sequences associated with coral thermal tolerance. The genes implicated by this analysis include (a) aspects of core cellular metabolism (e.g. transcription and protein metabolism) and also (b) known components of the transcriptional stress response in corals (e.g. heat shock proteins, reactive oxygen stress responses proteins). We hypothesize that this sequence variation contributes to thermal tolerance in corals by (a) minimizing the damage (thermal denaturation of proteins with roles in core cellular metabolism), and (b) enhancing the activity or stability of proteins that counter these effects (e.g. responses to unfolded proteins or reactive oxygen species).

Many of the genes and classes of genes differentially expressed in this study corroborate findings in other transcriptomic studies of other species of corals highlighting broad generalities in response to heat stress. Only two GO annotation terms were identified in enrichment analysis: protein folding (GO:0006457) and response to abiotic stress (GO:0006950). This class of genes has been here was an upregulation in expression of several heatshock proteins (HSP) in response to elevated temperature treatments. Orthologs to HSP90a and 90b were both upregulated during thermal stress as has been demonstrated in other scleractinian corals (e.g. DeSalvo *et al.*, 2008, Gates & Edmunds, 1999, Meyer *et al.*, 2011, Rosic *et al.*, 2014, Yuyama *et al.*, 2012). Likewise HSP70 and HSP20 orthologues were upregulated in our dataset indicating that the canonical heat stress response was observed in larvae exposed to elevated temperatures.

Recently, Ruiz-Jones and Palumbi (2017) hypothesized a model where the unfolded protein response (UPR), responsible in part for maintaining endoplasmic reticulum (ER) homeostasis, is triggered in corals during periods of elevated temperatures. They found enrichment for genes involved in calcium binding and homeostasis during periods of high temperature associated with the tidal cycle in the coral *Acropora hyacinthus* in American Samoa. Although calcium homeostasis genes were not enriched in our dataset, we found differential expression of several genes that were associated with this process. We saw upregulation of several calumenin-like proteins, which are calcium binding enzymes that aid in protein folding in the ER (Sahoo & Kim, 2008), when larvae were exposed to 36° C. These genes were also upregulated in response to increased temperature in acclimated *Acropora millepora* colonies, but was downregulated in non-acclimated replicates (Bellantuono *et al.*, 2012a). This indicates that the larvae in our elevated temperature treatment may have been more tolerant than naïve colonies of *A. millepora*. Upregulation of two additional genes (ERO1 and ERp29) in response to elevated temperature further supports the hypothesis that heat induces ER stress. Both of these proteins increase in abundance in response to cellular and chemical stress in mammalian systems (Li *et al.*, 2009, Mkrtchian *et al.*, 1998). Finally, although not significant after multiple test correction (p= 0.0004, BH p=0.165), the endoplasmic reticulum (GO:0005783) was one of the highest scoring categories in the cellular component GO annotations. Taken together, our data support the model of ER stress detailed in Ruiz-Jones and Palumbi (2017).

The largest fraction of differentially expressed genes occurred in the interaction term of temperature and sire indicating large transcriptional differences between the thermally tolerant and susceptible colonies. These included several reverse transcriptases (RT), integrases and recombinases that were all significantly downregulated in the offspring sired by the least thermally tolerant parent (64), and upregulated in the most thermally tolerant parents (53 & 58) (Figure 4c). Upregulation of several RTs were also seen in *Orbicella faveolata* fragments exposed to thermal accompanied by a decrease in expression of the putative regulatory gene in the PIWI-like subfamily (DeSalvo *et al.*, 2008). The functional consequences of this variation remain unclear, but the scale of these differences and their observation across multiple systems suggests they warrant further study.

Analysis of variation in expressed sequences revealed extensive genetic variation associated with thermal tolerance, like the multilocus SNP genotyping (2bRAD) described above. In this case the SNPs are directly associated with genes that can be functionally annotated, providing a rich source of information on the functional categories of genes differentiating heat-tolerant from heat-susceptible corals. We found extensive variation in gene sequences associated with coral thermal tolerance. The genes implicated by this analysis include (a) aspects of core cellular metabolism (e.g. transcription and protein metabolism) and also (b) known components of the transcriptional stress response in corals (e.g. heat shock proteins and reactive oxygen stress responses proteins). We hypothesize that this sequence variation contributes to thermal tolerance in corals by (a) minimizing the damage (thermal denaturation of proteins with roles in core cellular metabolism), and (b) enhancing the activity or stability of proteins that counter these effects (e.g. responses to unfolded proteins or reactive oxygen species).

Overall, our study demonstrates that despite strong selection for thermal tolerance in the Gulf, corals in this region retain substantial variation in thermal tolerance. Strong selection for thermal tolerance would be expected to deplete this variation, suggesting that something acts to maintain this variation in the face of ongoing selection. Further studies should clarify the relative roles of gene flow and balancing selection in maintaining this variation. This system has enormous potential for expanding our understanding of the thermal limits for coral adaptation. Ongoing efforts to develop genomic resources for this system will improve functional interpretations of genomic studies in this system, and provide new insights into the genes and processes underlying the unusual thermal tolerance of Gulf corals.

## Authors’ contributions

NK conducted crosses and larval experiments, helped with preparation and analysis of sequencing libraries, conducted heritability analysis, and helped write the manuscript. EJH and DA helped conduct crosses and larval experiments. JB provided crucial logistical support for the experiments in his laboratory. EM supervised all sequence analysis in his laboratory, prepared and analyzed sequencing libraries, and helped write the manuscript. All authors played a role in conceiving the study and all authors read and approved the final manuscript.

## Acknowledgements

The authors are grateful to Grace Vaughan and Dain McParland at New York University Abu Dhabi for their help with field work and aquarium set up. Permits for coral collection were provided by the Environment Agency Abu Dhabi (<permit information pending>.

## Supplementary file descriptions

Supplementary File S1. FASTA file. *De novo* reference produced by clustering 2bRAD sequences from aposymbiotic larval samples (SRA accession SUB3015376), used as a reference for 2bRAD analysis in this study.

Supplementary Table S2. Excel spreadsheet. Summary of sequencing yields, processing, and mapping efficiencies.

Supplementary Table S3. Excel spreadsheet. Complete list of SNPs significantly associated with thermal tolerance.

Supplementary Table S4. Excel spreadsheet. Complete table of differentially expressed genes.

Supplementary Table S5. Excel spreadsheet. Output of functional enrichment analysis on gene expression data.

Supplementary Table S6. Excel spreadsheet. Complete list of transcripts containing SNPs significantly associated with thermal tolerance.

